# 20-hydroxyecdysone (20E) primes innate immune responses that limit bacteria and malaria parasite survival in *Anopheles gambiae*

**DOI:** 10.1101/818526

**Authors:** Rebekah A. Reynolds, Hyeogsun Kwon, Ryan C. Smith

## Abstract

Blood-feeding is an integral behavior of mosquitoes to acquire nutritional resources needed for reproduction. This requirement also enables mosquitoes to serve as efficient vectors to acquire and potentially transmit a multitude of mosquito-borne diseases, most notably malaria. Recent studies suggest that mosquito immunity is stimulated following a blood meal, independent of infection status. Since blood-feeding results in the increased production of the hormone 20-hydroxyecdysone (20E), we hypothesized that 20E may play an important role in priming the immune response for pathogen challenge. Herein, we examine the immunological effects of priming in *Anopheles gambiae* with 20E prior to pathogen infection, demonstrating a significant reduction in bacteria and *Plasmodium berghei* survival in the mosquito host. RNA-seq analysis following 20E treatment identifies several known 20E-regulated genes, as well as several immune genes with previously reported function in anti-pathogen defense. This includes the anti-microbial peptide cecropin 3, which we demonstrate its role as an antagonist of bacteria and *Plasmodium* in *Anopheles gambiae* and provide support that these responses are under temporal regulation. Together, these data demonstrate that 20E influences cellular immune function and anti-pathogen immunity following mosquito blood-feeding, arguing the importance of hormones in the regulation of mosquito innate immune function.

## Introduction

Blood-feeding behavior evolved in mosquitoes to provide nutritional resources required for egg development. While providing benefits for reproduction, blood-feeding also exposes mosquitoes to a myriad of blood-borne pathogens that can ultimately be transmitted to a new host through an additional blood meal. For this reason, mosquitoes are arguably the deadliest animals on the planet, causing hundreds of millions of infections and over 500,000 deaths every year. Of mosquito-borne diseases, malaria continues to be the most deadly, where more than 400,000 people die annually from *Plasmodium* parasite infection transmitted by female *Anopheles* mosquitoes [1].

Following a blood meal, signals initiated in the brain produce ovarian ecdysteroidogenic hormone (OEH) and insulin-like peptides (ILPs), which trigger ecdysone production by the ovaries [2–4]. Ecdysone is then converted into 20-hydroxyecdysone (20E) by hydroxylation in the fat body, stimulating the production of yolk protein precursors (YPPs) in a process known as vitellogenesis [2,5–7]. Reaching peak levels approximately 18-24 hours after blood-feeding [8,9], 20E expression also coincides with *Plasmodium* ookinete invasion of the midgut epithelium [10]. While the influence of 20E is well-established with respect to insect development [11], mating [12,13], reproduction [7,14,15], and vectorial capacity [16,17], the role of 20E expression on mosquito immunity remains relatively unexplored.

Evidence from other insect systems suggests that 20E is an important regulator of host innate immune responses [18–22]. In *Drosophila*, 20E initiates immune signaling through the IMD pathway via regulation of the peptidoglycan receptor PGRP-LC, or through PGRP-LC-independent mechanisms, to directly regulate a subset of antimicrobial genes for anti-bacterial defense [20]. However, only a limited number of studies have addressed the influence of 20E on mosquito innate immunity [23,24]. Previous studies implicate 20E in prophenoloxidase (PPO) expression [23] and the regulation of leucine-rich repeat immune protein 9 (LRIM9) [24], both of which contribute to anti-*Plasmodium* immunity [24,25]. Additional studies also implicate 20E in mediating the effects of blood-feeding on mosquito immune cell activation and protein expression [26–28], suggesting that 20E is an important determinant of *An. gambiae* cellular immune function.

To better understand the role of 20E in mosquito immunity, we examine the ability of 20E herein to mediate phagocytic activity and influence immune priming prior to pathogen challenge. We demonstrate that 20E increases phagocytic activity, and reduces bacteria and *Plasmodium* survival. Through RNA-seq analysis, we show that 20E application induces known components of 20E signaling and several immune genes implicated in mosquito immunity, suggesting that 20E primes mosquito innate immunity for pathogen challenge.

## Methods

### Mosquito rearing and *Plasmodium* infections

A colony of *An. gambiae* G3 was maintained at 27°C and 80% relative humidity, with a 14/10 hour light/dark cycle. Mosquito larvae were fed on a ground fish food diet, while adult mosquitoes were maintained on a 10% sucrose solution.

For mosquito infections, a mCherry strain of *Plasmodium berghei* [29] was passaged into female Swiss Webster mice and monitored for exflagellation as previously described [25,30,31]. Mosquitoes were fed on an infected, anesthetized mouse, then maintained at 19°C. Mosquito midguts were dissected eight days post-infection in 1x PBS, and oocyst numbers were measured by fluorescence using a Nikon Eclipse 50i microscope.

### RNA-seq analysis

To examine the effects of 20E on mosquito cells, we applied 500ng of 20E (dissolved in 100% EtOH) to Sua 4.0 cells and harvested cells for RNA isolation 24 hours post-20E application. Sua 4.0 cells treated with a comparable volume of 100% EtOH for 24 hours were used as controls. Total RNA was isolated using TRIzol (Thermo Fisher Scientific), then purified with the RNA Clean & Concentrator-5 kit (Zymo Research) and quantified using a Nanodrop spectrophotometer (Thermo Fisher Scientific). RNA quality and integrity were measured using an Agilent 2100 Bioanalyzer Nano Chip (Agilent Technologies). 200 ng of total RNA was used to perform RNA-seq analysis from four independent biological replicates. RNA-seq libraries were prepared by the Iowa State University DNA Facility using the TruSeq Stranded mRNA Sample Prep Kit (Illumina) using dual indexing according to the manufacturer’s instructions. The size, quality, and concentration of the libraries were measured using an Agilent 2100 Bioanalyzer and a Qubit 4 Fluorometer (Invitrogen), then diluted to 2 nM based on the size and concentration of the stock libraries. Clustering of the libraries into a single lane of the flow cell was performed with an Illumina cBot. 150 base paired-end sequencing was performed on an Illumina HiSeq 3000 using standard protocols.

Raw sequencing data were analyzed by the Iowa State Genome Informatics Facility. Sequence quality was assessed using FastQC (v 0.11.5) [32], then paired-end reads were mapped to the *Anopheles gambiae* PEST reference genome (AgamP4.9) downloaded from VectorBase [33] using STAR aligner (v 2.5.2b) [34]. Genome indexing was performed using the genomeGenerate option, and corresponding GTF file downloaded from VectorBase (version 4.7) followed by mapping using the alignReads option. Output SAM files were sorted and converted to BAM format using SAMTools (v 1.3.1) [35], and counts for each gene feature were determined from these alignment files using featureCounts (v 1.5.1) [36]. Reads that were multi-mapped, chimeric, or fragments with missing ends were excluded. Counts for each sample were merged using AWK script, and differential gene expression analyses were performed using edgeR [37]. Differentially expressed genes with a q-score ≤ 0.1 were considered significant and were used for downstream analyses. Gene expression data have been deposited in NCBI’s Gene Expression Omnibus [38] and are accessible through GEO accession number GSE116252 (https://www.ncbi.nlm.nih.gov/geo/query/acc.cgi?acc=GSE116252).

### *In vivo* injection of 20E in mosquitoes

20E (Sigma) was resuspended in 138 uL of 100% EtOH and diluted in 1X PBS (1:10) for *in vivo* injections. 69nL of 20E (500ng) was injected into the thorax of anesthetized mosquitoes. Surviving mosquitoes were challenged 24 hours after 20E priming, or samples were collected for RNA isolation as previously described [24]. Control mosquitoes were injected with 69 nL of 100% EtOH diluted in 1X PBS (1:10).

### Gene expression analysis

Total RNA was isolated from Sua 4.0 cells or whole mosquitoes (~10 mosquitoes) using TRIzol (Thermo Fisher Scientific) according to the manufacturer’s protocol. RNA samples were quantified using a Nanodrop spectrophotometer (Thermo Fisher Scientific) and ~2ug of total RNA was used as a template for cDNA synthesis using the RevertAid First Strand cDNA Synthesis kit (Thermo Fisher). Gene expression was measured by qRT-PCR using gene-specific primers (Table S1) and PowerSYBR Green (Invitrogen). The results were normalized to an S7 reference gene and quantified using the 2^−ΔΔCT^ method as previously [39]. Samples were run in triplicate for each experiment.

### dsRNA synthesis and gene-silencing

T7 primers for GFP and *cecropin 3* (cec3) were used to amplify cDNA prepared from whole *An. gambiae* mosquitoes and cloned into a pJET1.2 vector using a CloneJET PCR Cloning Kit (Thermo Fisher). The resulting plasmids were used as a template for amplification using the corresponding T7 primers (Table S2). PCR products were purified using the DNA Clean and Concentrator kit (Zymo Research) and used as a template for dsRNA synthesis as previously described [30,31]. The resulting dsRNA was resuspended in RNase-free water to a concentration of 3 µg/uL. For gene-silencing experiments, 69nL of dsRNA was injected per mosquito and evaluated by qRT-PCR to establish gene knockdowns at 1-5 days post-injection. Time points with the highest efficiency of gene-silencing were chosen for downstream experiments.

### Phagocytosis Assays

*In vivo* phagocytosis assays were performed as previously described [25,31]. Briefly, mosquitoes were injected with 69nL of red fluorescent FluoSpheres (1 μm, Molecular Probes) at a 1:10 dilution in 1x PBS. Following injection, mosquitoes were allowed to recover for two hours at 19°C, then perfused onto a multi-test slide. Samples were fixed using 4% paraformaldehyde for 30 minutes, washed three times using 1X PBS, then blocked in 1% BSA for 30 minutes at room temperature. Hemocytes were stained using a 1:500 dilution of FITC-labeled wheat germ agglutinin (WGA; Sigma) in 1x PBS and incubated overnight at 4°C. After washing with 1x PBS, cell nuclei were stained with ProLong Gold anti-fade reagent with DAPI (Invitrogen). Hemocytes were identified by the presence of WGA and DAPI signals, and the percent phagocytosis was calculated by dividing the number of cells containing red fluorescent beads by the total number of cells present. The phagocytic index was calculated by counting the total number of beads per cell (this is summed for all of the cells) dividing by the number of phagocytic cells. Approximately 200 cells were counted per mosquito sample.

### *E. coli* challenge

To evaluate the effect of 20E priming on bacterial infection, challenge experiments were performed as previously [40] with control or 20E-treated mosquitoes. Briefly, kanamycin-resistant *E. coli* were cultured at 37°C until reaching an optical density (OD) of 0.4. Approximately 1mL of the *E. coli* solution was centrifuged at 10,000xg for 10 minutes, and the supernatant was removed. The pellet was washed twice with 1x PBS, then concentrated, before resuspending in 1x PBS. Challenge experiments were performed by injecting 69nL of *E.coli* into the thorax of each mosquito under each experimental condition. After injection, *E. coli*-challenged mosquitoes were kept at 19°C for 12 to 24 hours before perfusion. The hemolymph from five mosquitoes (~50uL) was pooled and diluted in 450uL of room temperature LB broth. 100uL of the pooled hemolymph sample was plated on LB agar plates containing kanamycin in triplicate and incubated overnight at 37°C. The number of *E. coli* colonies per plate were recorded, with the average used to compare between control and 20E-treated samples (number of *E. coli* colonies/average of control). This standardization was used to normalize for variation in *E. coli* numbers between experiments.

To determine the effects of *Cec3* gene-silencing on bacterial load, a slightly modified procedure was used to evaluate individual mosquitoes bacteria titers [41,42]. Briefly, control and *Cec3*-silenced mosquitoes were challenged two days post-dsRNA injection with 69nL of kanamycin-resistant *E. coli* as described above. 24 hours post-*E. coli* challenge, individual mosquitoes were homogenized in 100µL of 1xPBS. Samples were then diluted with an additional 100 µL of 1xPBS, vortexed, and 50 uL of each sample plated on kanamycin agar plates. Plates were incubated overnight at 37°C, and the number of colonies was counted and converted to colony forming units/mL.

## Results

### Blood-feeding and 20E injection increase phagocytic activity

To determine the effects of 20E on mosquito immune function, we first explored whether blood-feeding and 20E influenced mosquito cellular immunity. Immune cells, known as hemocytes, serve as primary immune sentinels that recognize and destroy invading pathogens by phagocytosis or the production of humoral immune factors [43,44]. To examine how blood-feeding and 20E influence these phagocytic properties, we injected fluorescent beads to evaluate phagocytosis under different physiological conditions as previously [25,31]. Similar to Kwon and Smith [25], we demonstrate that blood-feeding, independent of pathogen challenge, significantly increased the percentage of phagocytic cells (Figure 1A) and phagocytic activity (Figure 1B). We therefore hypothesized that the increase in phagocytic activity may be influenced in part by the increased levels of 20E post-blood-feeding. To address this question, we injected 20E into mosquitoes prior to challenge with fluorescent beads. Similarly, we observed a significant increase in the proportion of phagocytic cells (Figure 1C) and phagocytic activity (Figure 1D) 24 hours after 20E injection, suggesting that 20E activates mosquito cellular immunity.

**Figure 1.**
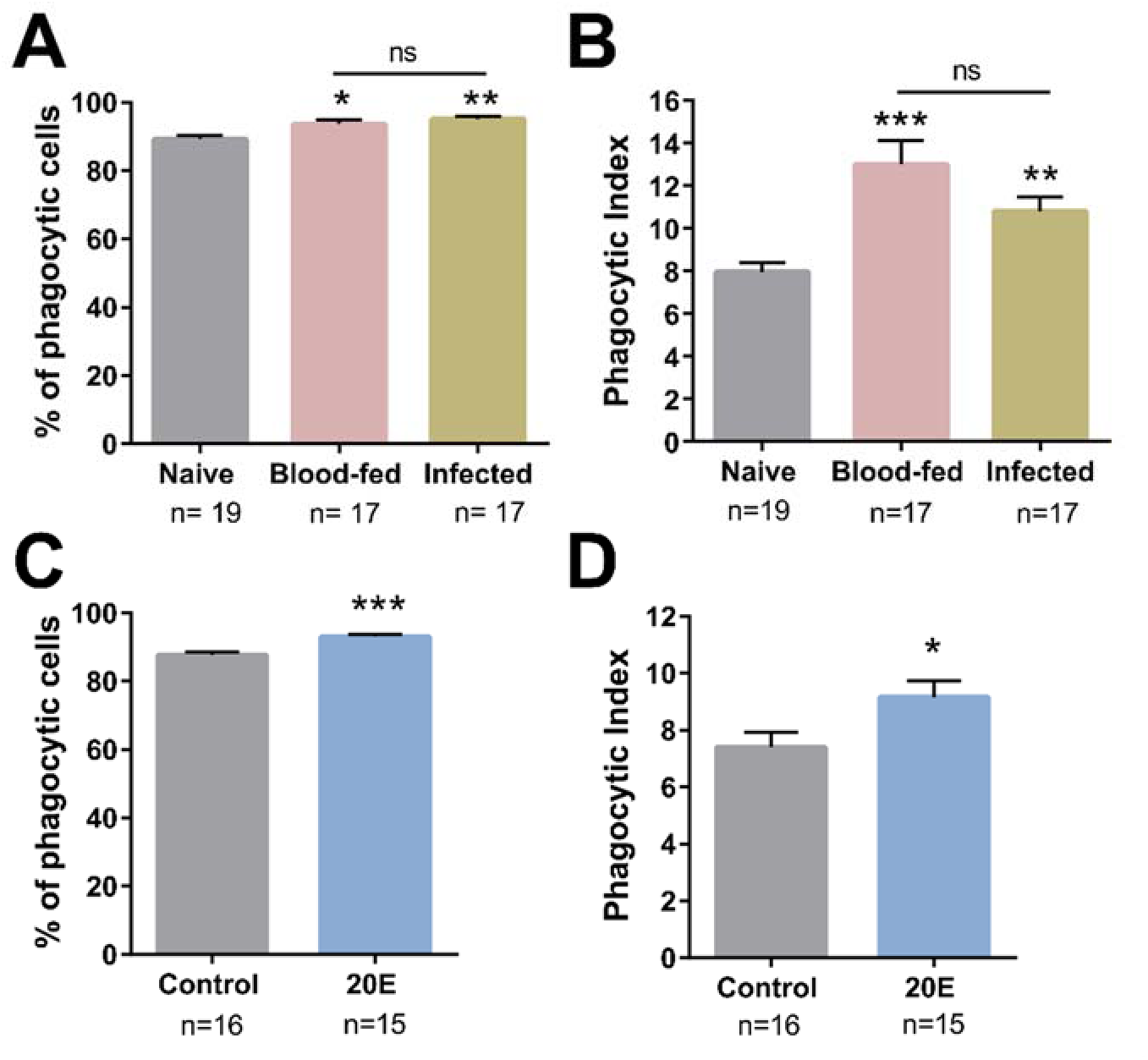
Phagocytic activity increases following blood feeding and 20E injection. Phagocytosis assays were performed in adult female *An. gambiae* under naïve, blood-fed, or *P. berghei*-infected conditions approximately 24 hours post-feeding. Perfused hemocytes from each condition were evaluated by the percentage of phagocytic cells (**A**) and the phagocytic index (number of beads per cell) (**B**). Similar experiments were performed following 20E priming, where the influence of 20E was evaluated by the percentage of phagocytic cells (**C**) and the phagocytic index (**D**). Three or more independent experiments were performed for each treatment. Data were analyzed by Kruskal-Wallis with a Dunn’s post-test (**A** and **B**) or Mann-Whitney (**C** and **D**). n= the number of mosquitoes examined for each condition. Asterisks denote significance (\**P* < 0.05, \*\**P* < 0.01, \*\*\**P* < 0.001); ns, not significant.

### Blood-feeding and 20E limit bacterial infection

Based on our observations that blood-feeding and 20E increase phagocytic activity, we next looked to determine the role of blood-feeding and 20E priming on bacterial challenge. To approach this question, we challenged naïve and blood-fed mosquitoes (~24 hours post-feeding) with *E. coli* and bacterial titers were evaluated 18 to 24 hours later from perfused hemolymph. Comparisons of naïve and blood-fed mosquitoes revealed a significant reduction in bacterial numbers in blood-fed mosquitoes when *E. coli* titers were examined at 24 hours (Figure 2A). Since 20E is expressed following a blood meal, we speculated 20E signaling increased immune function and subsequently reduced *E. coli* survival. This was examined by priming mosquitoes with 20E, then challenging as previously with *E.coli* at 12, 18, or 24 hours post-20E priming. We found a significant reduction in *E. coli* titers when challenged 12 hours post-20E priming (Figure 2B), yet the effects of 20E priming were abrogated when challenged at 18 (Figure 2C) or 24 hours post-20E priming (Figure 2D), suggesting 20E transiently primes anti-bacterial immunity. Interestingly, this peak effect on anti-bacterial immunity occurs ~12 hours post 20E-priming. This corresponds to the approximate timing of peak 20E production in the mosquito host, where 20E is induced ~12 hours post-blood meal and peaks between 18-24 hours post-blood feeding [45].

**Figure 2.**
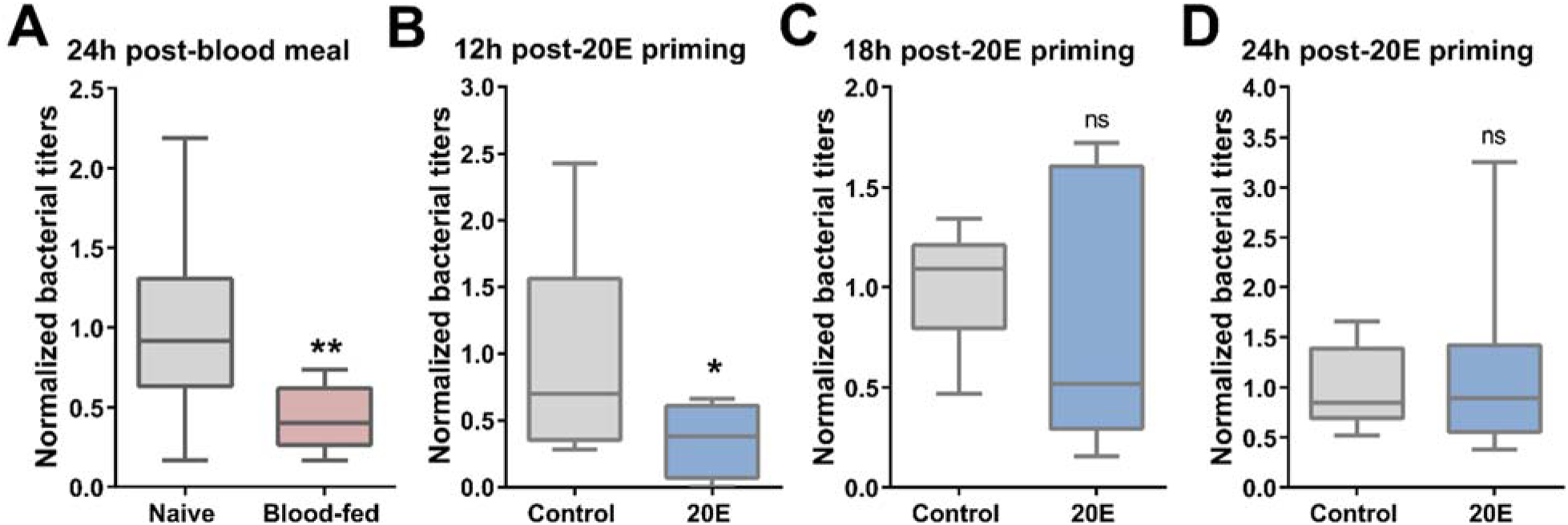
Blood-feeding and 20E priming reduce *E. coli* survival. The effects of blood-feeding and 20E priming on bacteria titers were examined through bacteria challenge experiments with *E. coli*. Mosquitoes were challenged with bacteria ~24 h post-blood meal (**A**), 12h post-20E priming (**B**), 18 h post-20E priming (**C**), or 24 h post-20E priming (**D**). For each experimental condition, hemolymph bacteria titers were examined 24 h later by perfusion. Pooled hemolymph from five mosquitoes was plated in duplicate on LB-KanR plates and the bacterial colonies were counted to determine titers. Data were normalized from three or more independent experiments and visualized using box plots (whiskers show min./max. values, line denotes median value) and evaluated using Mann-Whitney to determine significance (* = *P* < 0.05, ** = *P* < 0.01); ns, not significant.

### 20E priming limits *Plasmodium* survival

Since 20E influences phagocytosis and anti-bacterial immunity (Figures 1 and 2), we wanted to explore whether 20E priming similarly influences malaria parasite numbers in the mosquito host. To examine this question, we primed mosquitoes with 20E, then challenged with *P. berghei*. We found the injection of 20E 24 hours before *Plasmodium* infection significantly reduced parasite survival (Figure 3A) and the prevalence of infection (Figure 3B), suggesting that 20E initiates anti-*Plasmodium* immune responses that prime the mosquito host.

**Figure 3.**
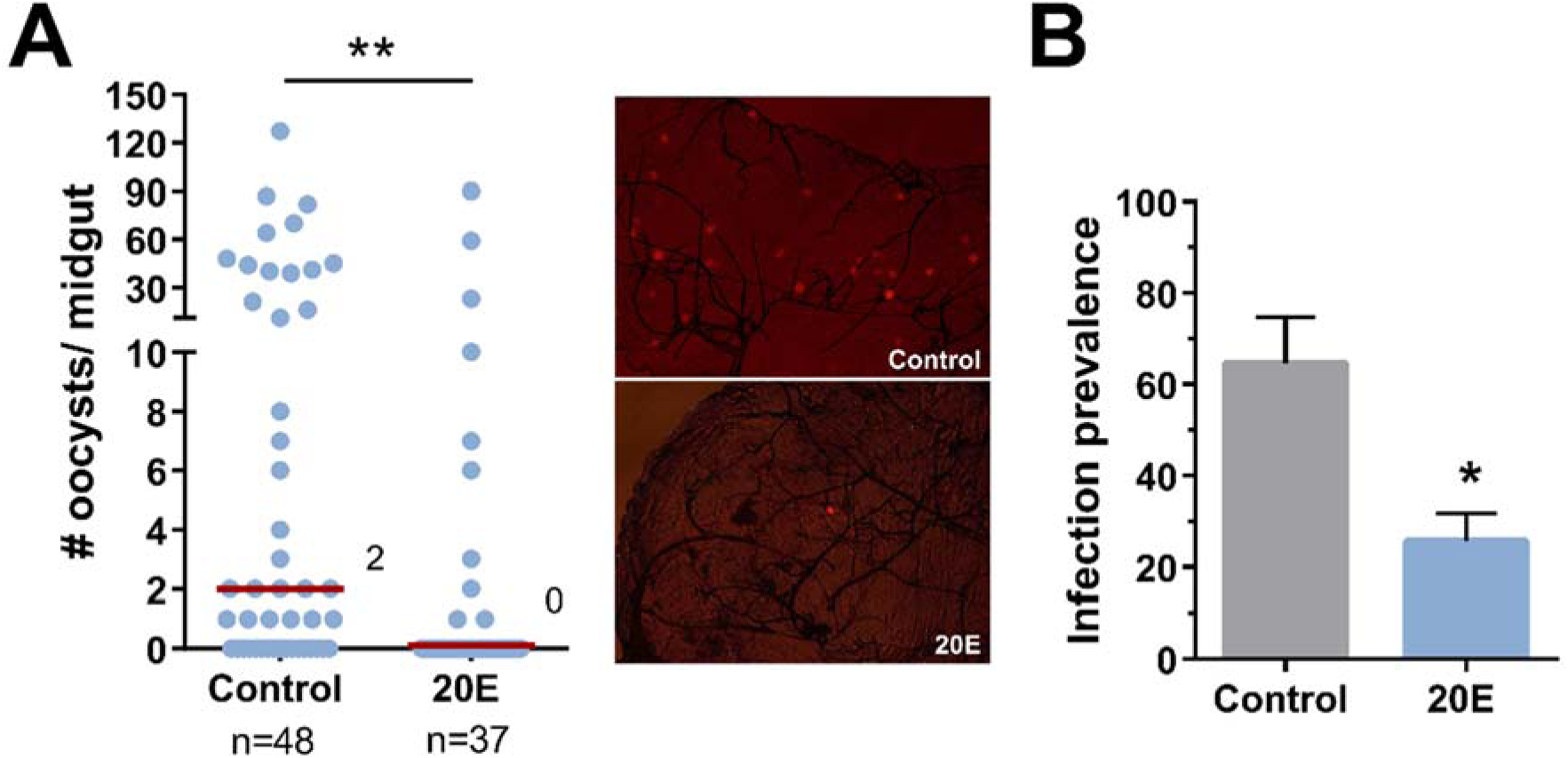
20E priming significantly reduces *Plasmodium* survival. Adult female mosquitoes were injected with 10% EtOH in 1xPBS (control) or 20E, then challenged with a *P. berghei*-infected mouse 24 h later. Eight days post-infection, *Plasmodium* oocyst numbers were evaluated by fluorescence as shown through representative images for each condition (**A**). Data were pooled from four independent experiments. Median oocyst numbers from both treatments are displayed by a red line. The percentage of mosquitoes containing at least one oocyst (prevalence of infection) was combined from each of the four indepent experiments (**B**). A Mann-Whitney was used to determine significance (* = *P* < 0.05, ** = *P* < 0.01); n= number of mosquitoes examined per treatment.

### 20E-regulation of mosquito immunity

To better understand how 20E primes immune responses to bacteria and malaria parasites, we applied 20E to *An. gambiae* Sua 4.0 cells to identify genes under the direct or indirect regulation of 20E. Our analysis identified 128 differentially regulated genes, including 80 up-regulated genes and 48 down-regulated genes (Table S3). Gene ontology analysis demonstrates that 20E application up-regulates transcripts associated with transport, immunity, and transcription, while down-regulating transcripts with predicted function in metabolism and redox metabolism (Figure 4A). This includes the upregulation of several previously described 20E-induced “early genes”, *HR3*, *HR4*, *E75A/B*, and multiple *broad* complex isoforms (Figure 4B, Table S3), implicated in canonical 20E signaling [46].

**Figure 4.**
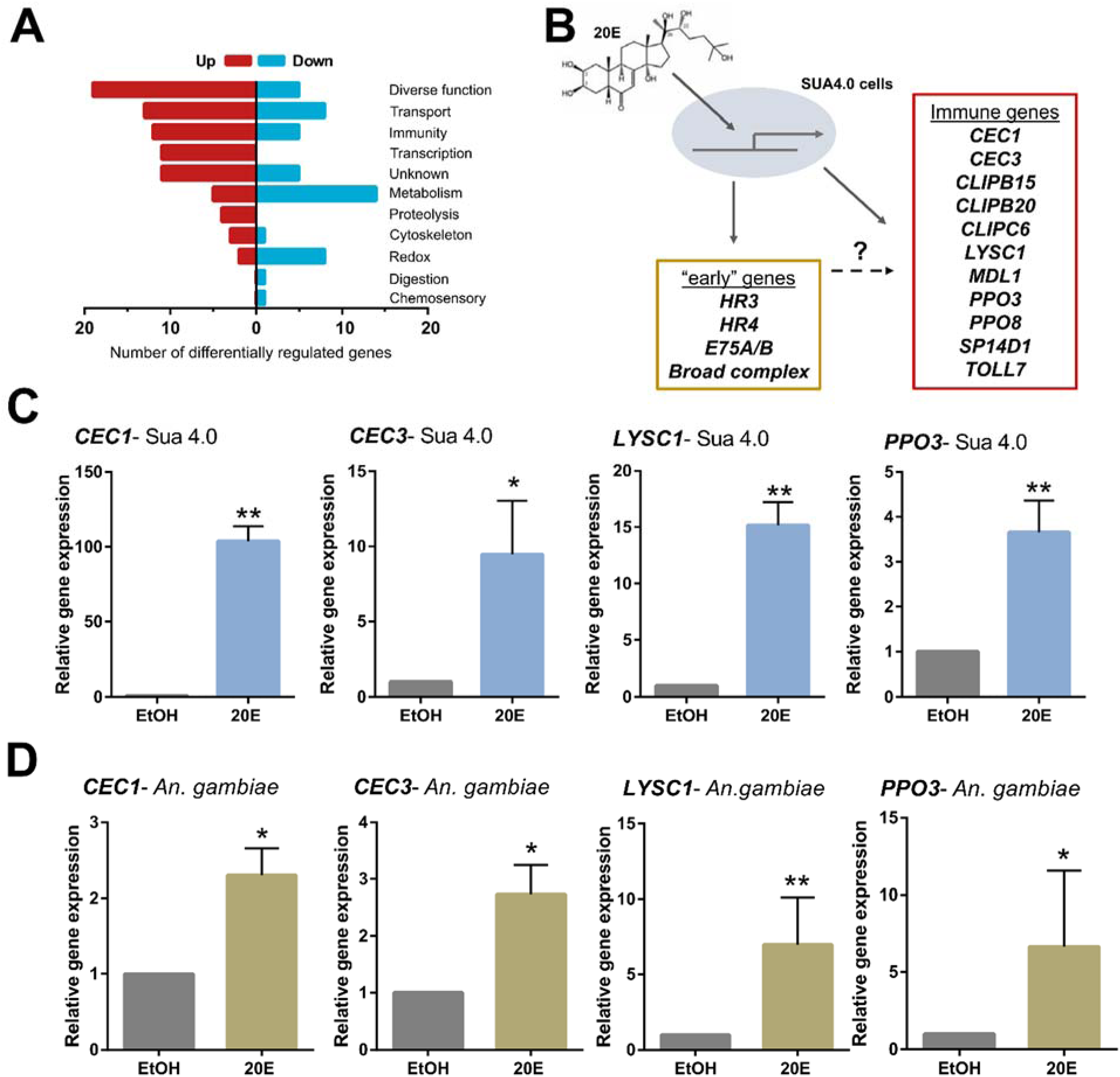
RNA sequencing identifies immune genes stimulated by 20E application. RNA-seq analyses of the effects of 20E on Sua 4.0 cells identified 128 differentially regulated genes (80 up-regulated, 48 down-regulated) grouped by gene ontology (**A**). Included among these genes were a number of previously described “early-genes” under 20E regulation, as well as multiple genes involved in innate immunity (**B**). Four of these immune genes (CEC1, CEC3, LYSC1, PPO3) were examined to validate the effects of 20E in either Sua 4.0 cells (**C**) or in vivo in whole female *An. gambiae* samples (**D**) compared to control (EtOH) treatments. Statistical significance was determined by Mann-Whitney analysis (* = *P* < 0.05, ** = *P* < 0.01) from four independent biological samples.

In addition, 20E application up-regulated 12 genes associated with mosquito immune function (Figure 4B, Table S1). PPO3 is a known *Plasmodium* antagonist [25], while three other genes, LYSC1, CLIPB15 and MDL1 are antagonists against both bacteria and malaria parasites [47–51]. The anti-microbial peptides, CEC1 and CEC3 (Figure 4B, Table S1), are widely implicated in general anti-pathogen effects to both bacteria and parasites [41,52,53], and while the exact function of CLIPC6, CLIPB20, and SP14D1 remains undetermined, researchers previously implicated these members of the CLIP protease family in mosquito immune signaling [54,55]. Together, this suggests 20E treatment profoundly affects mosquito immunity through the regulation of several previously characterized immune genes.

We validated our RNA-seq results using a subset of differentially regulated genes *in vitro* (Figure 4C, Figure S1) and *in vivo* using whole mosquito samples following 20E injection (Figure 4D, Figure S1) by qRT-PCR. Gene expression significantly correlated with both *in vitro* and *in vivo* samples (Figure S1), although gene expression results more closely matched the *in vitro* samples as expected. These differences are exemplified by differences in the expression of several genes between mosquito cell lines and whole mosquito samples where 20E only significantly influences expression *in vitro* (Figure S1). However, we cannot exclude that 20E may significantly influence the expression of these transcripts in specific mosquito tissues.

### Cecropin 3 (CEC3) limits bacterial and malaria parasite survival

Since several of the immune genes under 20E regulation are previously described, we further examined the role of CEC3 on *E. coli* and *Plasmodium* survival. Following the injection of dsRNA, we efficiently silenced CEC3 (Figure 5A), thus enabling experiments to determine the role of CEC3 in mediating, in part, the effects of 20E priming. To address CEC3 function on bacterial titers, we challenged individual control and *cec3*-silenced mosquitoes with *E. coli*. Under naïve conditions, *cec3*-silencing significantly increased bacterial titers, yet did not affect blood-fed mosquitoes (Figure 5B). This suggests CEC3 contributes to anti-bacterial responses under naïve conditions similar to previously described studies [41], yet a large number of immune components induced by blood-feeding which contribute to anti-bacterial immunity mask the influence of CEC3 on bacterial titers [24,26,28].

**Figure 5.**
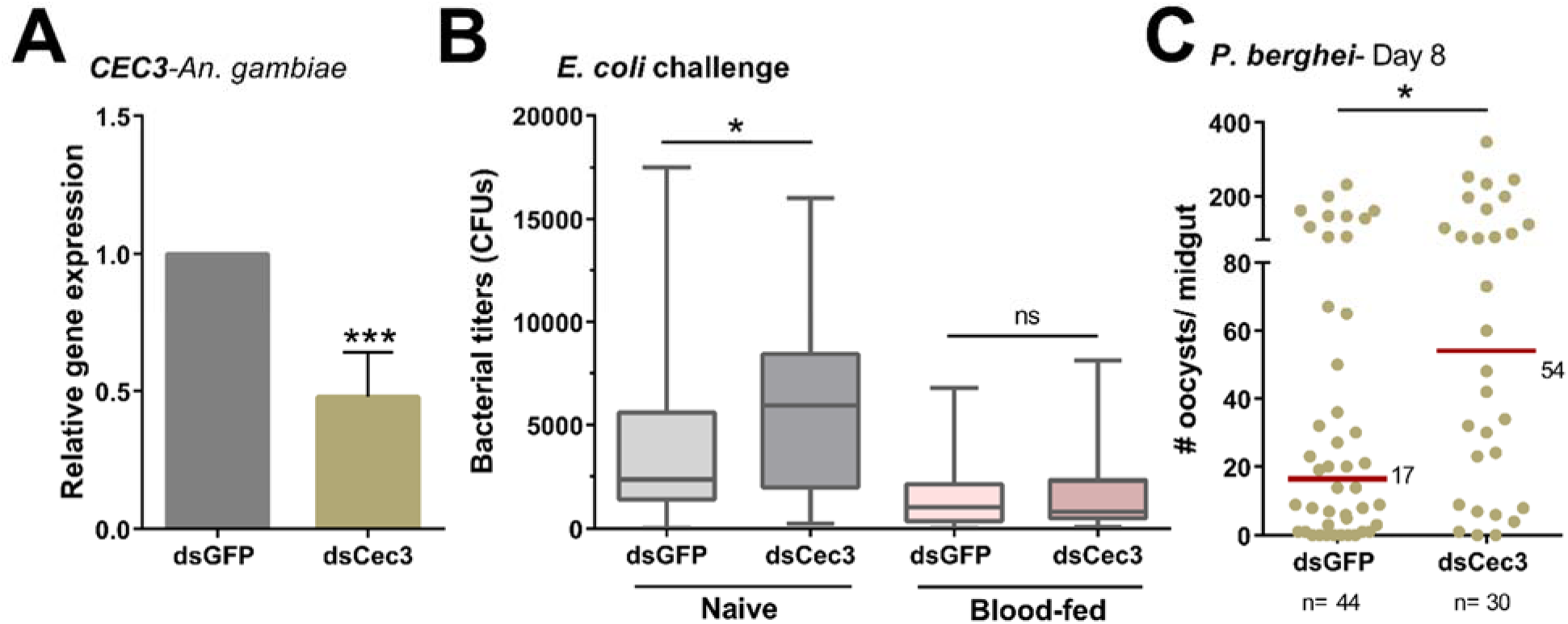
*Cecropin 3* impacts antibacterial and anti-*Plasmodium* immunity. The efficiency of *cec3*-silencing was examined in whole mosquitoes by qRT-PCR by comparing *cec3* transcript in dsGFP (control) or dsCec3 samples (**A**). Statistical analysis was performed using a students t-test from four independent experiments. To determine the role of *cec3* in reponse to bacterial challenge, bacterial titers were examined in individual mosquitoes following the injection of dsGFP (control) or dsCec3 under naïve and blood-fed conditions (**B**). More than thirty mosquitoes were examined for each experimental condition, with data analyzed using a Mann-Whitney test to determine significance. In addition, the effects of cec3 silencing were examined on P. berghei oocyst numbers eight days post-infection and analyzed using Mann-Whitney (**C**). For each experiment asterisks denote significance (* = *P* < 0.05, *** = *P* < 0.001); ns, not significant; n= number of mosquitoes examined per treatment.

We also investigated the role of CEC3 on *P. berghei* infection. Compared to control mosquitoes, *cec3*-silencing resulted in a significant increase in oocyst numbers (Figure 5C), suggesting CEC3 also contributes to anti-*Plasmodium* immunity similar to previous studies [41,52,53]. Together, these experiments demonstrate that CEC3 provides generalized anti-pathogen effects that mediate both bacterial and malaria parasite survival.

## Discussion

Blood-feeding is an essential behavior for *An. gambiae* and other hematophagous insects to acquire nutrient resources for egg production. This requires the synthesis of yolk protein precursors in a process known as vitellogenesis that is regulated by the production of 20E shortly after a blood-meal [7,15,56]. In addition, the male transfer of 20E during mating further contributes to oogenesis to maximize female fecundity [12,13,57]. However, studies of 20E function on the immune system of mosquitoes and other insects thus far are limited.

Much of our current understanding of 20E function in innate immunity relies on previous studies in *Drosophila* [19–22]. 20E signaling regulates anti-microbial protein (AMP) production in *Drosophila* [19,20] through PGRP-LC-dependent and -independent mechanisms via the ecdysone receptor and ultraspiracle [20]. Evidence form *Drosophila* also suggests that 20E promotes hemocyte activation, leading to increased mobility, responsiveness to wounding, and phagocytic activity [22,58]. Through the results presented herein, we see striking similarities in which 20E regulates cecropins (AMPs) and increased phagocytic activity. We also find that 20E mediates anti-pathogen effects on both bacteria and malaria parasites (Figure 6), similar to the anti-bacterial effects of 20E signaling described in *Drosophila* [20,22]. Despite these parallels, there are distinct physiological differences between *Drosophila* and *An. gambiae*, most notably in blood-feeding behavior of mosquitoes, pathogen exposure, and the production of 20E. For this reason, there is reason to believe the role of 20E may vary between *Drosophila* and mosquitoes, particularly regarding innate immunity.

**Figure 6.**
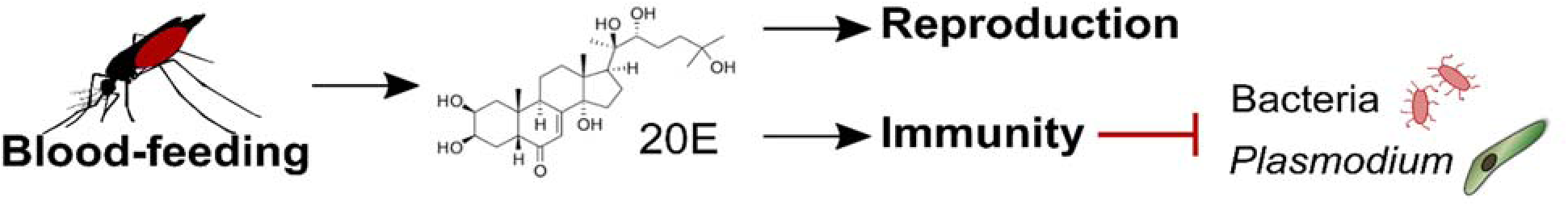
Multi-modal effects of blood-feeding and 20E on mosquito reproduction and immunity. Blood-feeding stimulates the production of 20E that can initiate vitellogeneis and influence mosquito fecundity, as well as prime mosquito immune responses that limit bacteria and malaria parasite survival.

In mosquitoes, several studies suggest that blood-feeding, independent of pathogen challenge, stimulate mosquito immunity [24,26–28]. However, these studies only indirectly implicate the role of 20E function in the mosquito immune response. More direct evidence demonstrates the role of 20E in prophenoloxidase (PPO) expression *in vitro* [23] and that of LRIM9, a leucine-rich immunomodulatory protein, through experiments *in vivo* [24]. Previous studies show both LRIM9 [24] and multiple PPOs [25] influence *Plasmodium* killing, supporting a model in which 20E promotes an anticipatory immune response to immune challenge immediately following a blood meal [24]. When combined with the anti-pathogen effects of CEC3 presented herein, and the previously described anti-bacterial and anti-*Plasmodium* responses of other immune genes identified in our RNA-seq analysis [47–51], our experiments demonstrate the widespread activation of mosquito innate immunity by 20E.

Given the role of 20E priming on bacteria and malaria parasites, 20E likely triggers both cellular and humoral immune components that limit pathogen survival. We demonstrate that 20E increases the phagocytic activity of mosquito immune cells, the production of AMPs, and the production of other immune elicitors, yet the precise mechanisms and tissues involved in these 20E immune priming responses remain unknown. In addition to changes in cellular immunity, humoral components produced by hemocytes or the fat body and secreted into the hemolymph may also act on bacteria or invading malaria parasites. Interestingly, our data suggest 20E priming differs temporally between pathogens, potentially providing additional resolution into the mechanisms of 20E-regulated immunity. The effects of 20E on *E. coli* appear more transient, with peak activity 12 to 18 hours post-priming. This contrasts with the longer-lasting influence of 20E immune priming on anti-*Plasmodium* immunity, where *P. berghei* challenge occurred 24 hours post-priming and persisted more than 18 hours post-infection after the onset of ookinete invasion. One potential explanation is that anti-bacterial responses are primarily cellular-mediated, while the immune responses that limit parasite development are humoral. This is supported by the transient activation of mosquito hemocytes following blood-feeding [26,27], which juvenile hormone (JH) may quickly suppress as described in *Drosophila* [19]. This is in contrast to protein-based humoral components, such as TEP1, which are stable in circulation in the mosquito hemolymph where they are integral to the recognition and lysis of invading ookinetes [10].

The establishment that 20E regulates mosquito immune responses raises additional considerations into the potential trade-offs between reproduction and immunity. Both physiological processes are energetically costly, where 20E production may serve as a limiting factor for the allocation of resources towards egg production or immunity [59]. Evidence suggests reduced mosquito fitness at the cost of anti-*Plasmodium* immunity [60,61], arguing a competition for resources in the mosquito host. This is supported by fitness experiments in transgenic mosquito lines expressing anti-*Plasmodium* effector genes which display increased fecundity compared to wild-type following *Plasmodium* challenge, presumably by enabling more resources for egg production [62,63]. However, recent studies argue *P. falciparum* infection intensity positively correlates with increased egg production and levels of 20E, where the production of lipids during vitellogenesis are used by *P. falciparum* to increase survival and optimize transmission [17]. Therefore, the effects of 20E on mosquito immunity and reproductive fitness potentially depend on specific host-pathogen interactions that define *An. gambiae* immunity to *P. berghei* and *P. falciparum* parasites [49,64] that require further study.

In summary, our findings demonstrate the role of the hormone, 20E, in priming mosquito innate immunity to both bacteria and malaria parasites. These effects are mediated in part through an activation of cellular immunity and likely involve a number of humoral factors, including that of cecropin 3 described herein, that make mosquitoes more resistant to pathogen infection. As a result, these data provide novel insights into the hormonal regulation of the mosquito immune system, yet require further investigation to better understand the regulation and tissue-specific contributions of 20E immune priming. Together with previous work [16], our results support that increasing 20E signaling in the mosquito host can reduce *Plasmodium* infection and may serve as a potential target to interrupt the transmission of malaria.

## Acknowledgements

BEI Resources, NIAID, NIH provided the *Anopheles gambiae* cell line: Sua 4.0 (MRA-921), which George Christophides initially contributed. National Science Foundation Graduate Research Fellowship Program under Grant No. 1744592 to R.A.R supported the work for this study. Additionally, the Agricultural Experiment Station at Iowa State University provided support for this research.

