## Supplementary material for "20-hydroxyecdysone (20E) primes innate immune responses that limit bacteria and malaria parasite survival in *Anopheles gambiae*": Figure S1, Table S1, Table S2

### Supplemental Information

### Supplemental Figures

**Figure S1.** Validation of RNA-seq data in Sua 4.0 cells and whole mosquito samples.

### Supplemental Tables

**Supplemental Table S1.** Primers used for qRt-PCR analysis.

**Supplemental Table S2.** Primers used for dsRNA synthesis.

**Supplemental Table S3.** Differentially regulated genes in Sua 4.0 cells following 20E treatment.

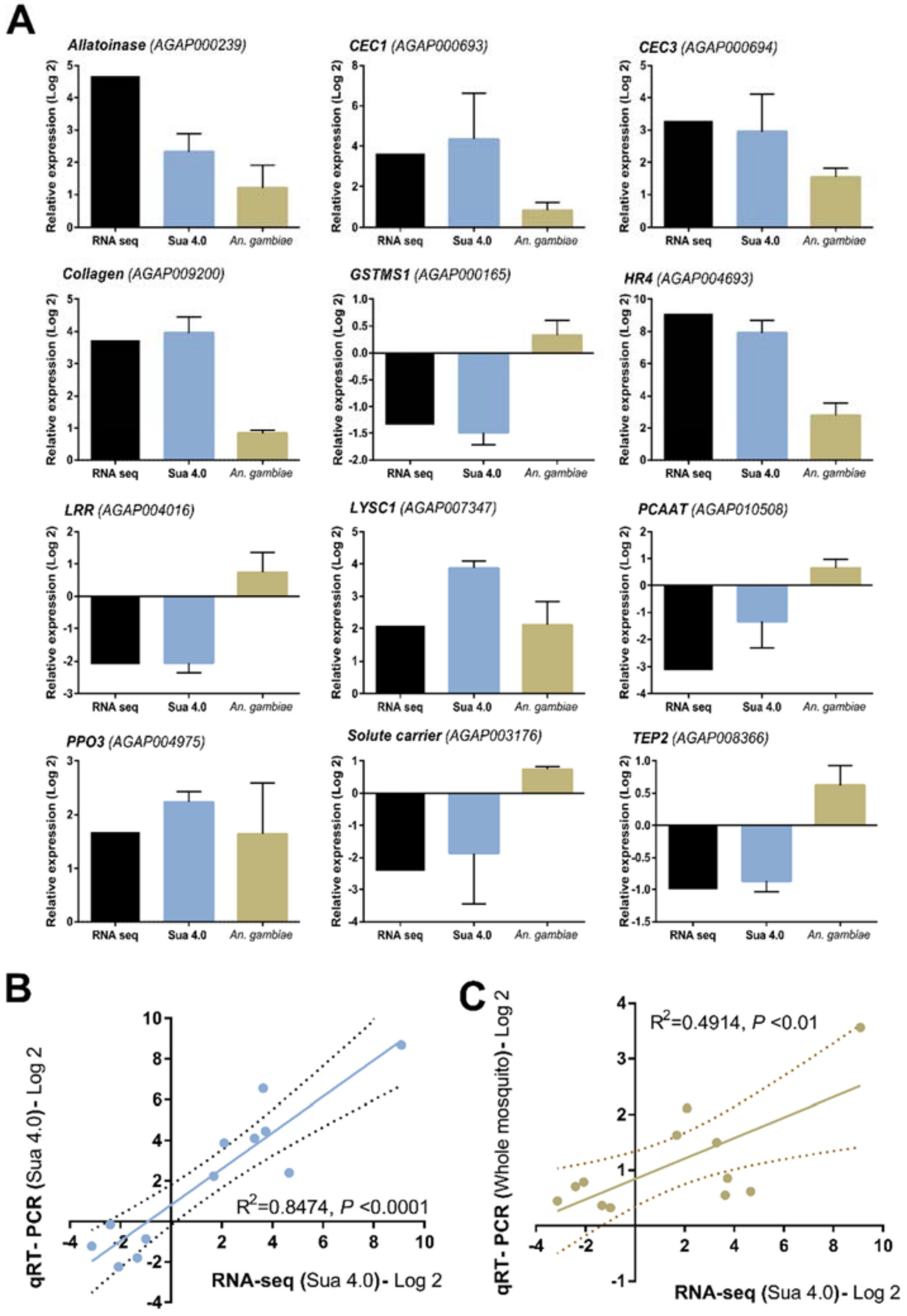

**Figure S1. Validation of RNA-seq data in Sua 4.0 cells and whole mosquito samples.** Differentially expressed genes identified in our RNA-seq analysis were validated independently in *An. gambiae* Sua 4.0 mosquito cells and whole *An. gambiae* female mosquitoes (G3 strain) by qRT-PCR (**A**). Expression data are displayed as the mean $\log_2$ fold change (±SEM) from at least three independent experiments. The RNA-seq data were correlated to qRT-PCR expression values using either Sua 4.0 cell samples (**B**) or whole mosquitoes (**C**) to validate the differential gene expression. To determine the significance, data was analyzed by a linear regression.

**Table S1. Primers used for qRT-PCR analysis**

| Primer | Gene ID | Sequence (5'- 3') |
|---|---|---|
| Allantoinase-F | AGAP000239 | CCCACGACGGTAAGGTGACG |
| Allantoinase-R | | TCCCGGCATGAGCATCAAAT |
| Cecropin 1-F | AGAP000693 | TTCATCTTTGTCGTGCTGGC |
| Cecropin 1-R | | GCACTGCCAGCACGACAAAG |
| Cecropin 3-F | AGAP000694 | ACGTACTGAACCACCTGCGCGTT |
| Cecropin 3-R | | GCGCTGTGTGCGCCGATGAA |
| Collagen-F | AGAP009200 | TGGGACTGCGAGGATTCGAG |
| Collagen-R | | ACCTCGTCCGACTGGCTGTG |
| GTMS1-F | AGAP000165 | GCGGTGCTGGTGGTGAAGAT |
| GTMS1-R | | CGTCCGGATCGTCGAACTTG |
| HR4-F | AGAP004693 | TCGGGGTCAAATGCATCACA |
| HR4-R | | GCTGCAGTTGGGTTCCGAGA |
| LRR-F | AGAP004016 | GCTCGTTTTGTGCCGGAATG |
| LRR-R | | GCCCGCTTCCTGCAGCTTAT |
| LYSC1-F | AGAP007347 | TTCAGCACATCGGCGACAAA |
| LYSC1-R | | TCCAGCCGTACCAGGCGTTA |
| PCAAT-F | AGAP010508 | TGCGGTGTTCCGGATATTGG |
| PCAAT-R | | AGGCGCCACTTCATCCATCC |
| PPO3-F | AGAP004975 | CTATTCGCCATGATCTCCAACTACG |
| PPO3-R | | ATGACAGTGTTGGTTGGTGAAACGGATCT |
| rpS7-F | AGAP010592 | ACCCCATCGAACACAAAGTTGACACT |
| rpS7-R | | CTCCGATCTTTCACATTCCAGTAGCAC |
| Solute Carrier-F | AGAP003176 | GTGCTTGGCTGTGTGCTGGA |
| Solute Carrier-R | | CGTTGGCCTGTACCGTCTCG |
| TEP2-F | AGAP008366 | GCACCTGGCTGACAGCGTTT |
| TEP2-R | | CCCTGACCCTGCACCTCCTT |

**Table S2. Primers used for dsRNA synthesis**

| Primer | Gene ID | Sequence (5'- 3') |
|---|---|---|
| GFP-T7-F | | TAATACGACTCACTATAGGGAGAATGGTGAGCAAGGGCGAGGAGCTGT |
| GFP-T7-R | | TAATACGACTCACTATAGGGAGATTACTTGTACAGCTCGTCCATGCC |
| Cecropin 3-T7-F | AGAP000694 | TAATACGACTCACTATAGGGTCAGTCTGAGATCTCTTCCCGT |
| Cecropin 3-T7-R | | TAATACGACTCACTATAGGGAGATTACTTGTACAGCTCGTCCATGCC |
